# Multi-plasmid clash in a bacterial community: plasmid viability depends on the ecological setting of hosts

**DOI:** 10.1101/2021.08.02.454727

**Authors:** Cindy Given, Reetta Penttinen, Matti Jalasvuori

## Abstract

Plasmids are genetic elements that disperse horizontally between different strains and species of bacteria and a major factor in the dissemination of virulence factors and antibiotic resistance. Understanding the ecology of plasmids has a notable anthropocentric value and therefore the interactions between bacterial hosts and individual plasmids have been studied in detail. However, bacterial systems often carry multiple genetically distinct plasmids, but dynamics of these multiplasmid “clashes” has remained unstudied. Here, we set to investigate the survival of 11 mobilizable or conjugative plasmids in five different ecological settings. The key incentive was to determine whether plasmid dynamics are reproducible and whether there are trade-offs in plasmid fitness that stem from the ecological situation of their initial hosts. Growth rates and maximum population densities increased in all communities and treatments over the 42-day evolution experiment although plasmid contents at the end varied notably. We show that large multiresistance conferring plasmids are unfit when the community also contains smaller plasmids with fewer resistance genes. This suggests that restraining the use to few antibiotics can make bacterial communities sensitive to others. The hosts also appear to react to the presence of multiple genetically different plasmids by enhancing fimbriae production instead of alleviating costs of individual plasmids. In general, the survivors of the here-studied multi-plasmid clash are significantly affected by the presence or absence of antibiotic selection and plasmid-free hosts of varying fitness. Therefore, these trade-offs in different settings can explain for example why some resistance plasmids have an advantage during a rapid proliferation of antibiotic sensitive pathogen whereas others dominate in alternative situations.

## Introduction

Bacterial cells serve as vehicles for various types of genetic replicators. In addition to chromosomes, these include transposing elements, plasmids, conjugative elements and viruses. Many of these replicators contain genes that can entirely dictate whether the hosting cellular vehicle (host) prevails in its present environment. For instance, replicators such as plasmids can contain antibiotic resistance genes that protect the host from drugs whereas viruses may directly destroy its host in order to spread the virus to the surrounding community (Jalasvuori and Koonin, 2015; Cairns et al., 2018). Both examples play a role in the antibiotic resistance crisis as plasmids disseminate the resistance phenotype and viruses have been championed to provide a cure for these antibiotic resistant bacteria (Furfaro et al., 2018; Decano et al., 2020). Overall, genetic replicators provide a framework for dissecting bacterial communities not only into cells of particular species with defined genetic contents but also into an interacting network of genetic entities (Jalasvuori and Koonin, 2015) and, arguably, evolutionary ecology of bacterial systems is also ecology of genetic replicators (Jalasvuori, 2012).

Plasmids are common genetic molecules of bacteria that (usually) replicate separately from the host chromosome (Norman et al., 2009). Plasmids are of notable anthropocentric importance due to their clinically relevant features, mainly antibiotic resistance and virulence factors (Lopatkin et al., 2017; Pilla and Tang, 2018). The presence of a (conjugative) plasmid in a particular strain can therefore both translate into the development of a disease and treatment failure. Conjugative plasmids encode a machinery that facilitates the transfer of the element into surrounding bacterial cells (Virolle et al., 2020). Mobilizable plasmid utilize these machineries to hitchhike along the way and disperse from one cell to another (Smillie et al., 2010). Among gram-negative pathogens, conjugative and mobilizable plasmids are the most common genetic replicators mediating multi-drug resistance (Carattoli, 2009; Partridge et al., 2018; Ruotsalainen et al., 2020).

Bacteria carry various types of plasmids that can be divided into incompatibility groups according to their capability to stably coexist in a single cell line. The general prevalence of plasmids has been puzzling to explain in the absence of selection for plasmid-carried genes, since in new hosts, plasmids often induce fitness costs (Carroll and Wong, 2018). These costs, however, may be rapidly alleviated by adaptive mutations (Dahlberg and Chao, 2003; Loftie-Eaton et al., 2017; Carroll and Wong, 2018). Conjugation rates also differ and the plasmid-associated costs to host fitness may also be balanced by their ability to invade surrounding cells (Lopatkin et al., 2017). In certain conditions such as under active predation by protozoa, plasmid survival appears to be dependent on their ability to conjugate (Cairns et al., 2016). Adaptation to specific plasmids can also render bacteria to become more permissive to other plasmids and the evolved plasmids, respectively, to be of lesser burden in alternative hosts (Harrison and Brockhurst, 2012; Loftie-Eaton et al., 2017). Different plasmids that have adapted to hosts of different bacterial species can come in contact later on and form successful novel plasmid combinations (Jordt et al., 2020; Alonso-Del Valle et al., 2021) and hence form new multi-resistant strains. Altogether, these studies draw a picture in which several factors contribute to the means by which plasmids and their hosts can form a stable long-term companionship. These factors together may resolve the so-called plasmid-paradox – in other words, explain why plasmids do not disappear when we withdraw the selection for specific plasmid-carried genes (as discussed by Carroll and Wong, 2018).

In reality, however, bacterial cells are often in an environment where multiple plasmids are continuously present and new plasmids migrate in within new hosts. Extended spectrum betalactamase producing plasmids identified from hospital settings are substantially diverse in their specific genetic characteristics as well as in their incompatibility groups and mobility types (Carattoli, 2009; Weingarten et al., 2018). Therefore, bacteria face situations where a variety of plasmids clash together to compete within host cells of close proximity (Bedhomme et al., 2017). Many basic questions in this domain remain mostly unstudied. Are there, for example, characteristics that make certain plasmids outcompete others during a multi-plasmid clash, and if so, how stochastic are these dynamics? Are there specific adaptations in the host in a situation where multiple genetically different plasmids occupy the community in comparison to often-studied situations where adaptations can focus on alleviating costs of singular plasmids (Bedhomme et al., 2017)? Is there always a single winner in a certain set of plasmids, or can the winner be dependent on the ecological setting of the host? Indeed, new plasmid-free bacterial hosts may either form within the community as some hosts lose plasmids or they may migrate in from other environments. These new migrants may also have higher fitness than members of the original community, and hence dispersal to new hosts can become essential for survival. To our knowledge, no studies have investigated the dynamics of such multi-plasmid clashes with more than three different plasmids and rarely in alternating ecological conditions. As such, exploring the answers to these questions served as an incentive for this study.

Here we set to investigate a set of plasmids with varying characteristics in differing ecological settings. Total of eleven different plasmids originating from clinical isolates were transferred to isogenic background (*Escherichia coli*) in their natural combinations and then subjected together to a 42-day long serial co-culture experiment in five different ecological settings (such as in the presence of migrating bacteria and antibiotics). The results show that some plasmids prevail in specific conditions while disappear in others. The host adaptation may be a response to several genetically different plasmids instead of being a specific response to one or a few. Overall, there appears to be trade-offs that may play a role in the success of some plasmids, e.g., in the presence of hosts with higher fitness.

## Results

We set up a serial culture microcosm experiment where each system was seeded with 11 plasmids that originated from extended-spectrum beta-lactamase (ESBL) encoding *Escherichia coli* strains in addition to a well-characterized conjugative plasmid RP4. The plasmids were transferred in their natural combinations to laboratory *E. coli* K-12 strain HMS174 in order to even out the competitive advantages that the original hosts. Some of the studied plasmids are conjugative and some are mobilizable (indicating that they are transferred along with a conjugative plasmid). In all plasmid combinations, at least one of the plasmids encode a beta-lactamase that provides resistance to betalactam antibiotics. Plasmid RP4 was included as it has been utilized in various studies where RP4-encoded conjugation machinery has been employed to deliver CRISPR-systems to antibiotic resistant bacteria (Bikard et al., 2014; Ruotsalainen et al., 2019). As such, RP4’s capability to compete amongst ESBL-plasmids provides estimates for its potential to modify bacterial communities with any introduced CRISPR-tools. However, RP4 contained a kanamycin resistance gene, which due to the experimental design, had to be inactivated prior to the microcosm experiments. Plasmids and their key features are listed in Table 1.

**Table 1.**
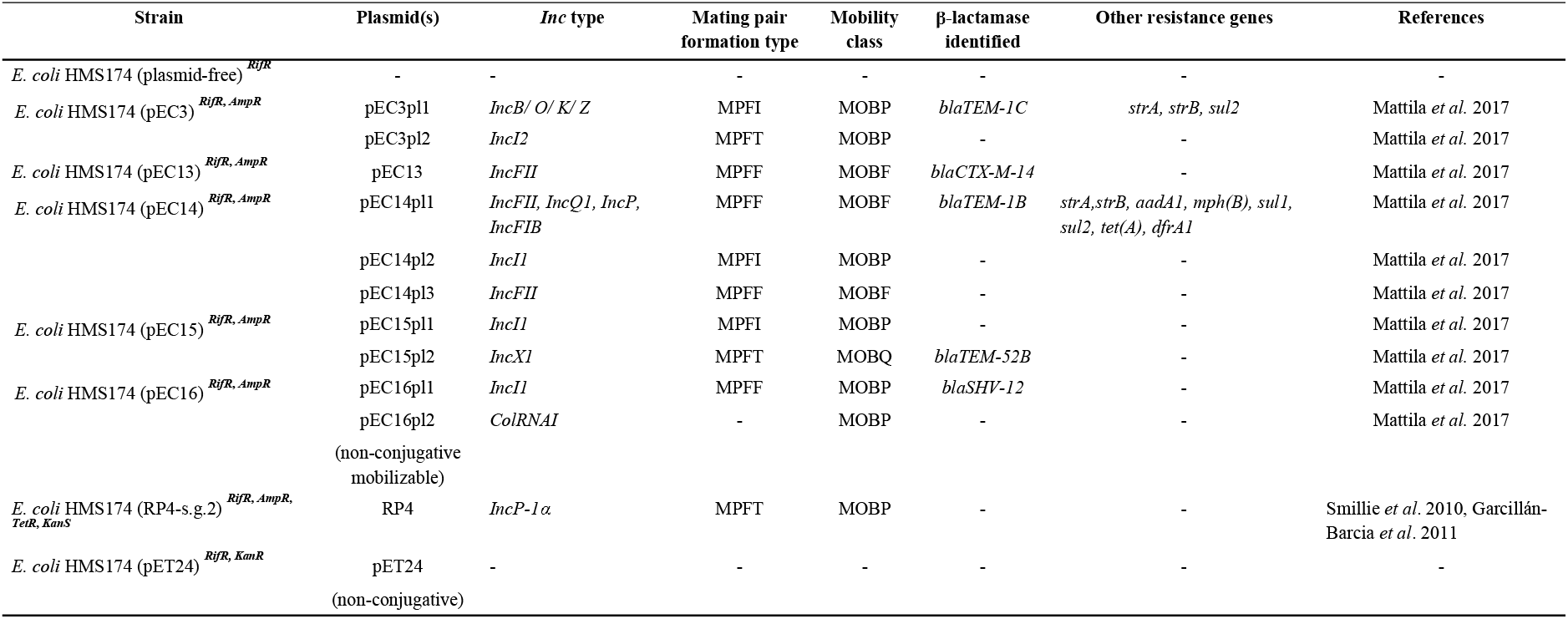
Plasmid-harboring strains used in the experiment

The plasmid-harbouring bacterial strains were mixed together for the microcosms that modelled five different ecological conditions (Figure 1). Each microcosm was refreshed 41 times, resulting in roughly 320 bacterial generations. Each experimental condition was carried out in four independent replicates. In the first setup, bacteria were cultivated and refreshed as a community (referred with C as in community). In the second setup, the community (CA, community + ampicillin) was subjected to continuous beta-lactam selection (thus representing an environment during treatment). In the third setup (CM, community + migration), bacterial community is continuously supplemented with plasmid-free hosts that can be invaded by the plasmids. In the fourth setup (CMK, community + migration + kanamycin), community is continuously introduced with migrating bacteria that have higher fitness than the original plasmid-hosts in an attempt to model the presence of rapidly proliferating plasmid-free pathogen. In the fifth setup (CMKA, community + migration + kanamycin + ampicillin), the community is treated with an antibiotic in the presence of an antibiotic-susceptible pathogen which could restore its full replication rate by receiving a resistance plasmid from the resistant community.

**Figure 1.**
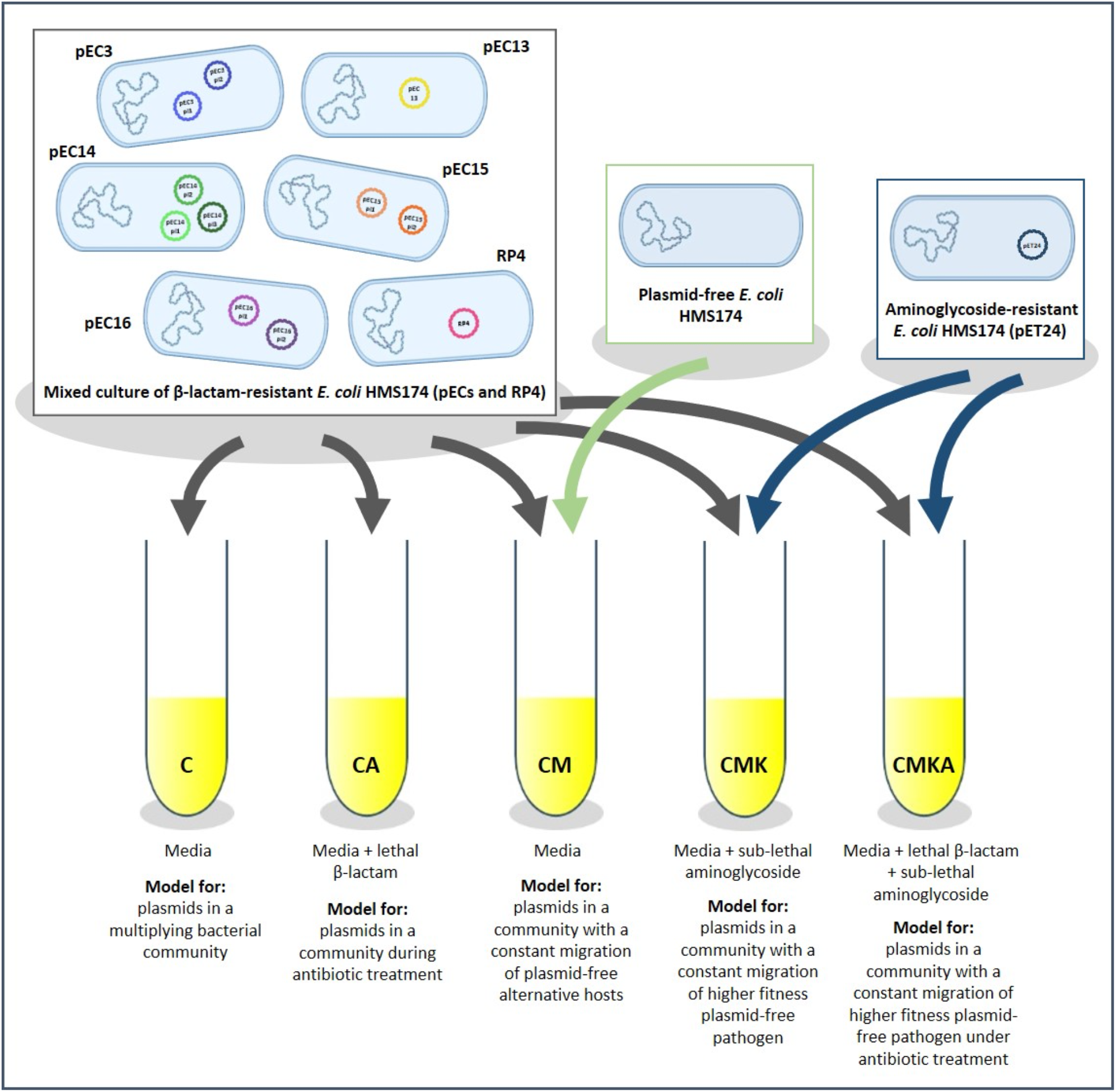
The experimental design of the serial culture experiment. Ampicillin was used for betalactam and kanamycin for aminoglycoside. The cultures were refreshed 41 times (N=4/treatment).

We measured the growth rates and maximum population densities for individual plasmid-hosting strains as well as for the evolved communities (Figure 2). All plasmids had non-significant effects on the replication rate of the hosts (Figure 2A). During the course of the experiment, the growth rate of the community increased significantly in all conditions (Figure 2B). Interestingly, the presence of plasmids increased the maximum population densities (yield) (Figure 2C). Evolved communities also showed increase in yield compared to the starting point (Figure 2D). Kanamycin concentration utilized in the experiment was selected for the kanamycin-resistant strain, HMS174(pET24), to have both higher replication rate and density compared to other strains in order to model the presence of higher-fitness plasmid-free host. We also measured the conjugation rates for all plasmid-harbouring hosts by their ability to transfer the beta-lactamase resistance to a susceptible recipient (Table 2). While this does not reveal the transfer rates for non-resistance plasmids, previous study with these plasmids indicated that all plasmids were co-transferred together (Mattila et al., 2016).

**Figure 2.**
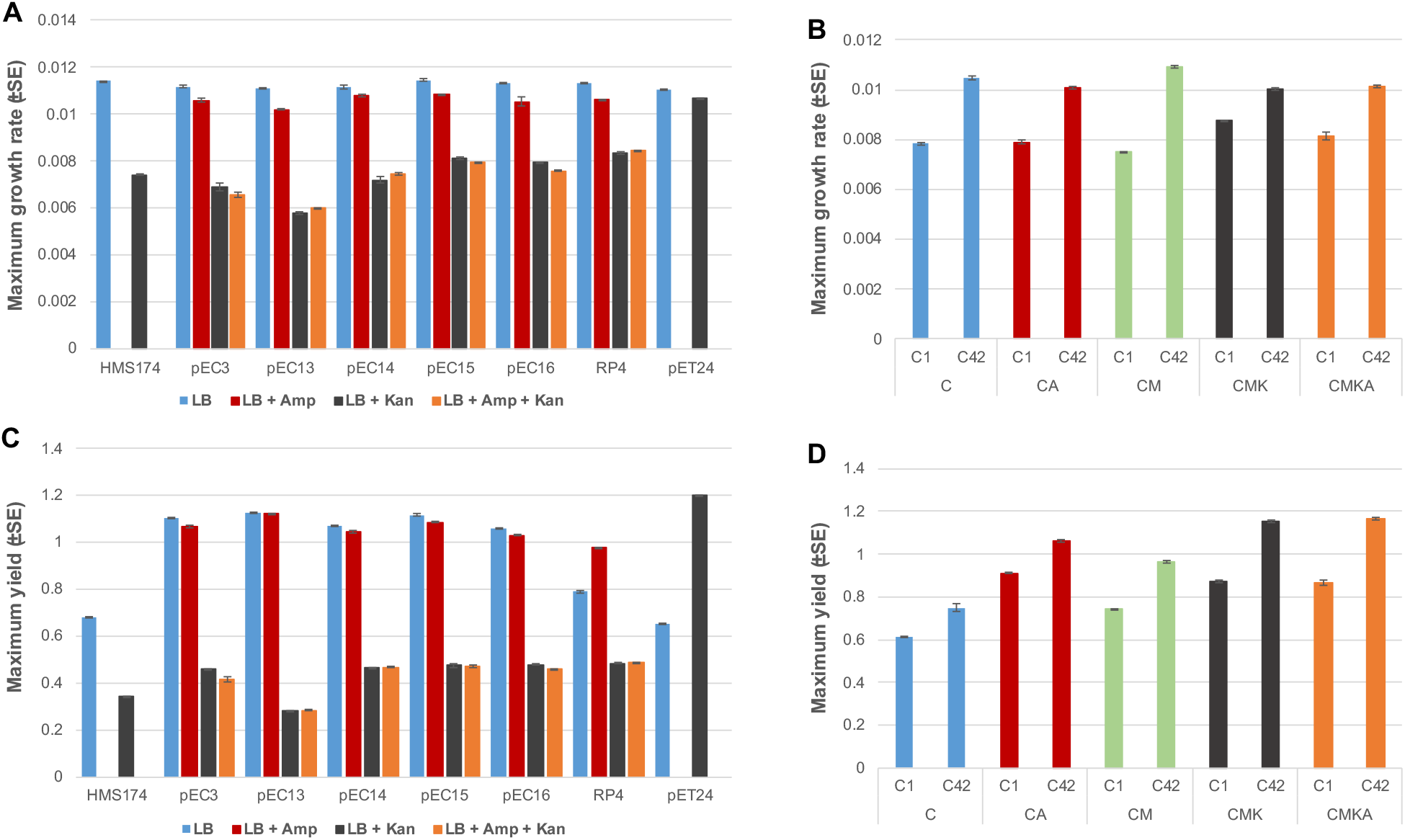
Maximum growth rates and yields of each individual strain (A and C) and communities under different treatments at the starting and end-point of the experiment (B and D). Statistical difference of growth rates and yields was determined using two-way ANOVA (p < 0.05). Growth rate for the CMK communities at starting point was the highest (p < 0.01, 2-way ANOVA, post-hoc: Tukey HSD) whereas no difference in the growth rate was shown between C, CA, and CMKA (p = 0.067, 2-way ANOVA, post-hoc: Tukey HSD). At the end-point, significant difference was observed among all treatments (p < 0.01, 2-way ANOVA, post-hoc: Tukey HSD) (B). Yields of the communities at the end point were significantly higher than the starting point between all treatments (p < 0.01, 2-way ANOVA, post-hoc: Tukey HSD) (D).

**Table 2:**
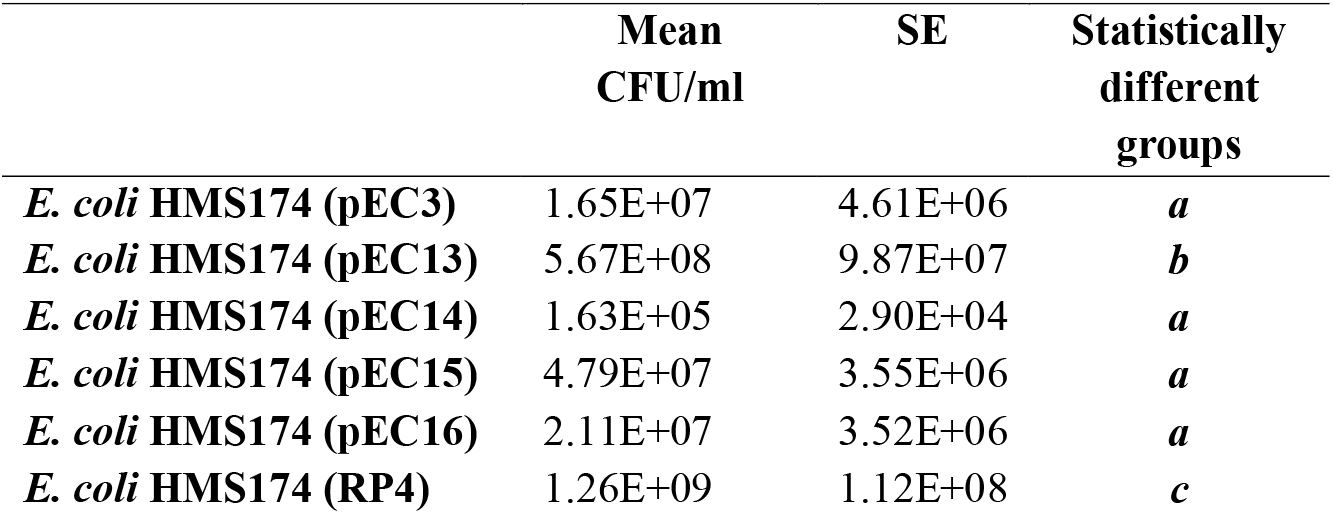
Average conjugation rate of beta-lactam resistance from each strain. Statistically significant differences are shown in bold italic letters behind the values (Two-way ANOVA, p<0.01).

Tracking the abundance of multiple plasmids with no specific selective genes can be challenging. As such, we employed quantitative polymerase chain reaction (qPCR) with specific primers against each plasmid (excluding pEC14pl2 and pEC15pl1 that were almost identical and hence were amplified with a single primer pair). Primer sequences and primer analysis are presented in Supplementary data 1. Total DNA was isolated at five time-points (1, 14, 32, 35 and 42 days) during the serial culture experiment and the relative quantity of each plasmid was determined against 16S rRNA genetic marker. qPCR measurements for each plasmid were conducted in three replicates. Figures 3 and 4 represent the evolution of plasmid prevalence over time (see Supplementary data 1 for original data). The majority of the eleven plasmids behaved similarly in all independent replicates within a particular experimental setup. However, pEC13 disappeared in all but one CA-replicate where it was well-represented, and pEC14pl3 disappeared in two CA-replicates and in one CM-replicate. This nevertheless suggests that plasmid-dynamics are generally consistent in a given ecological setting. Table 3 summarizes the behaviour of each plasmid in different setups.

**Figure 3.**
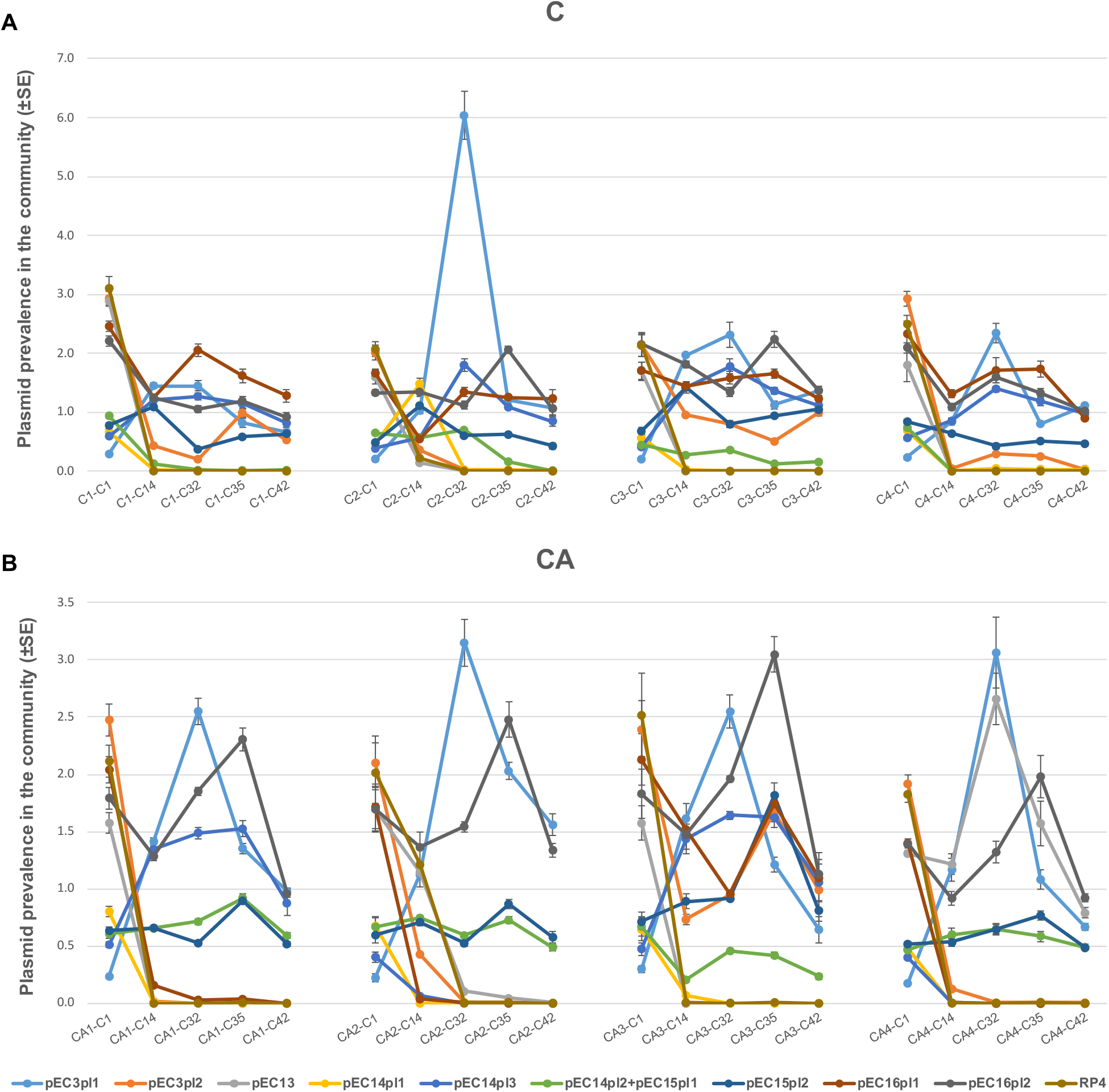
Prevalence of plasmids under different conditions as normalized with the gene for 16S ribosomal RNA over 42 cycles of the serial culture experiment. A) Four replicates of C “community”; B) Four replicates of CA “community with antibiotics” (beta-lactam ampicillin).

**Figure 4.**
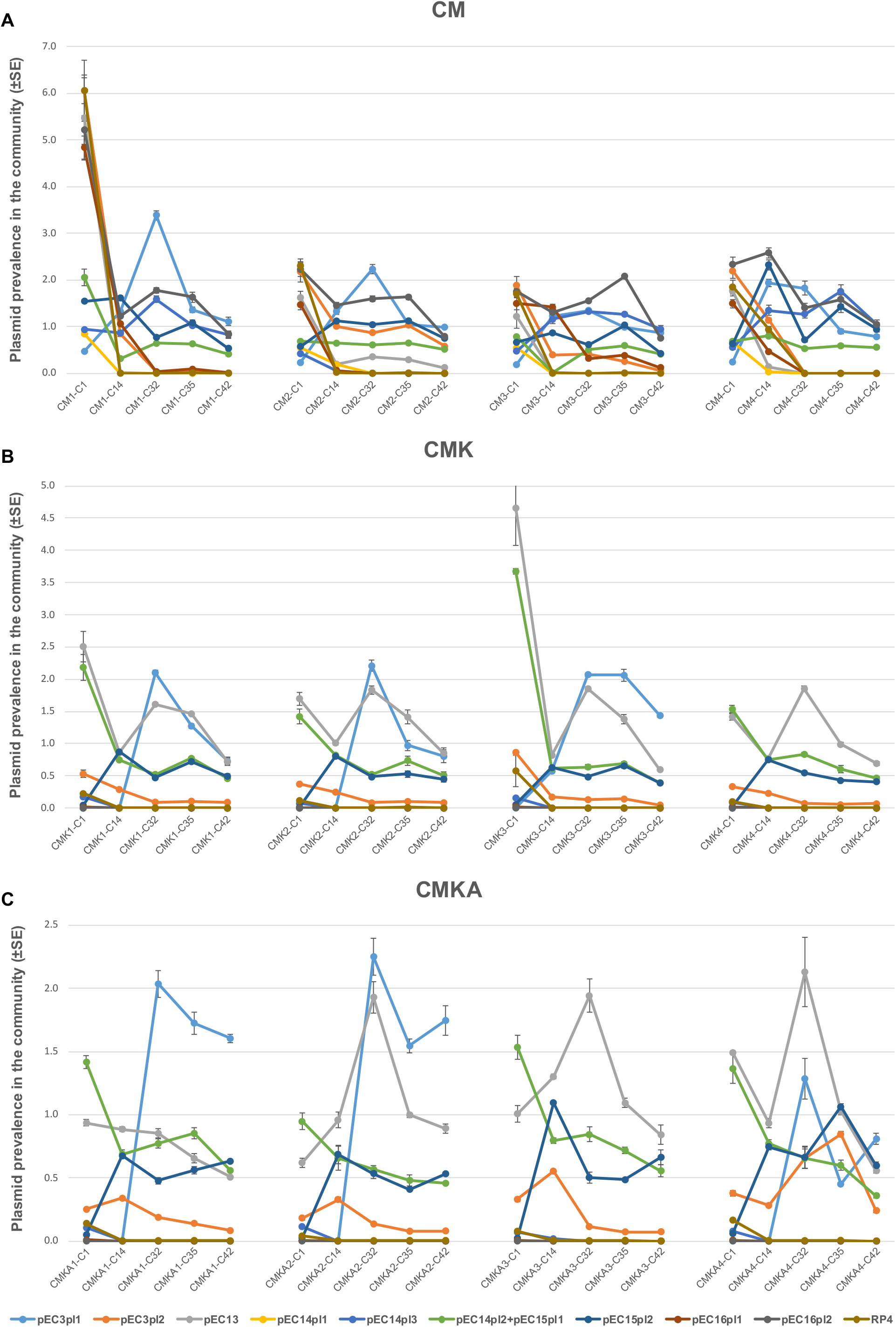
Prevalence of plasmids under different conditions as normalized with the gene for 16S ribosomal RNA over 42 cycles of the serial culture experiment. A) Four replicates of CM “community with migration of plasmid-free hosts”; B) Four replicates of CMK “community with migration of high-fitness plasmid-free hosts”; C) Four replicates of CMKA “community with antibiotics and migration of high-fitness plasmid-free (antibiotic sensitive) hosts”.

**Table 3.** Summary of plasmid dynamics in different ecological settings.

| Treatment | pEC3pl1 | pEC3pl2 | pEC13 | pEC14pl1 | pEC14pl3 | pEC14pl2+<br>pEC15pl1 | pEC15pl2 | pEC16pl1 | pEC16pl2 | RP4 |
| --- | --- | --- | --- | --- | --- | --- | --- | --- | --- | --- |
| **C** | Increase then decrease | Decrease and disappear | Decrease and disappear | Decrease and disappear | Increase then decrease | Decrease and disappear | Persist | Persist | Persist | Decrease and disappear |
| **CA** | Increase then decrease | Decrease and disappear | Decrease and disappear | Decrease and disappear | Increase then decrease /Decrease and disappear | Persist | Persist | Decrease and disappear | Persist | Decrease and disappear |
| **CM** | Increase then decrease | Decrease and disappear | Decrease and disappear | Decrease and disappear | Increase then decrease | Persist | Persist | Decrease and disappear | Persist | Decrease and disappear |
| **CMK** | Increase then decrease | Persist | Persist | Decrease and disappear | Decrease and disappear | Persist | Persist | Decrease and disappear | Decrease and disappear | Decrease and disappear |
| **CMKA** | Increase then decrease | Persist | Increase then decrease | Decrease and disappear | Decrease and disappear | Persist | Persist | Decrease and disappear | Decrease and disappear | Decrease and disappear |

We took a closer look at general features of plasmids in each of the ecological conditions to reveal characteristics that may be advantageous for plasmid survival (Table 4). Mobilizable plasmids can be grouped with MOB-classification system that bases its groups on relaxase genes (Garcillán-Barcia et al., 2009). Relaxase itself is responsible for both initiating and terminating conjugative transfer of a plasmid. Conjugative plasmids can also be classified by the protein complex that mediates the mating pair formation (MPF) with a recipient cell (Schröder and Lanka, 2005). In C, most of the plasmids with MOBP persisted in the system while all except one plasmid with MOBF disappeared (p = 0.001, 2-way ANOVA, post-hoc: Tukey HSD). In CA, plasmids with MOBP were more abundant than MOBF (p = 0.001, 2-way ANOVA, post-hoc: Tukey HSD). In CA, all plasmids with MPFI persisted in the system while most or half of plasmids with MPFT and MPFF disappeared from the system (p ≤ 0.01, 2-way ANOVA, post-hoc: Tukey HSD). In CM, all plasmids with MPFI persisted in the system while plasmids with MPFT and MPFF generally decreased and disappeared from the system (p ≤ 0.01, 2-way ANOVA, post-hoc: Tukey HSD). In CM, all except one of plasmids with MOBF were lost from the system while most plasmids with MOBP persisted (p = 0.005, 2-way ANOVA, post-hoc: Tukey HSD). In CMK, all plasmids with MPFI and MPFT persisted in the system while most of plasmids (except pEC13) with MPFF disappeared, however, statistically, MPFI was more abundant than MPFF (p = 0.001, 2-way ANOVA, post-hoc: Tukey HSD) and MPFT (p ≤ 0.01) but there was no difference between MPFF and MPFT (p = 0.621). In CMKA, all plasmids with MPFI and MPFT persisted in the system while most of the plasmids (except pEC13) with MPFF disappeared from the system (p ≤ 0.01, 2-way ANOVA, post-hoc: Tukey HSD). Here, MPFI was significantly more abundant than MPFF (p ≤ 0.01) and MPFT (p ≤ 0.01), but there was no difference between MPFT and MPFF (p = 1.000). In CMKA, no significant difference was observed among mobility groups (p = 0.142, 2-way ANOVA, post-hoc: Tukey HSD). It must be noted that, while there are potentially relevant benefits in different MOB and MPF-systems in different conditions, the plasmids vary in multiple characteristics and hence these results should be approached very cautiously.

**Table 4.** Persistence of different mobility and mating pair formation classes.

| Treatment | Mobility |  | p-value | Mating pair formation |  |  | p-value |
| --- | --- | --- | --- | --- | --- | --- | --- |
|  | MOBP | MOBF |  | MPFI | MPFT | MPFF |  |
| **C** | Most plasmids persisted | All but one plasmid disappeared | 0.001 | Most plasmids persisted | Most plasmids persisted | Most plasmids persisted | 0.919 |
| **CA** | Most plasmids persisted | Most plasmids decreased and disappeared | 0.001 | All plasmids persisted | Most plasmids disappeared | Half of the plasmids disappeared | ≤0.01 |
| **CM** | Most plasmids persisted | All but one plasmid disappeared | 0.005 | All plasmids persisted | Most plasmids decreased and disappeared | Most plasmids decreased and disappeared | ≤0.01 |
| **CMK** | Most plasmids persisted | One out of three plasmids persisted | 0.226 | All plasmids persisted | All plasmids persisted | Most of the plasmids disappeared | ≤0.01 |
| **CMKA** | Most plasmids persisted | Most plasmids persisted | 0.142 | All plasmids persisted | All plasmids persisted | Most plasmids disappeared | ≤0.01 |

We also investigated potential evolutionary changes in the plasmids. Sequencing of the end-point communities revealed some mutations that repeatedly appeared during the serial culture experiment. Most of the mutations in plasmids were located in non-coding regions or genes with unknown functions, excluding pEC3pl1 in which mutations within the gene for conjugal transfer protein V (TraV) appeared in multiple communities (Figure 5). Genetic changes in non-coding regions were close to transposases or shufflon regions that are known to be less stable due to continuous DNA “shuffling” (Brouwer et al., 2019). Bacterial host consistently accumulated mutations in genes for type 1 fimbriae regulatory protein (FimE) and in certain conditions (CM) a single nucleotide variant in gene for RecA (Figure 6). The low number of mutations in CMKA community is likely to derive from the continuous supplement of non-evolved HMS174 that had higher fitness compared to plasmid-harbouring strains. This produced constant migration of naïve mutation-free host that possessed advantageous non-transmissible plasmid pET24. No mutations in four conjugative plasmids (pEC13, pEC14pl3, pEC16pl2 and RP4) and pET24 were detected in any of replicate communities.

**Figure 5.**
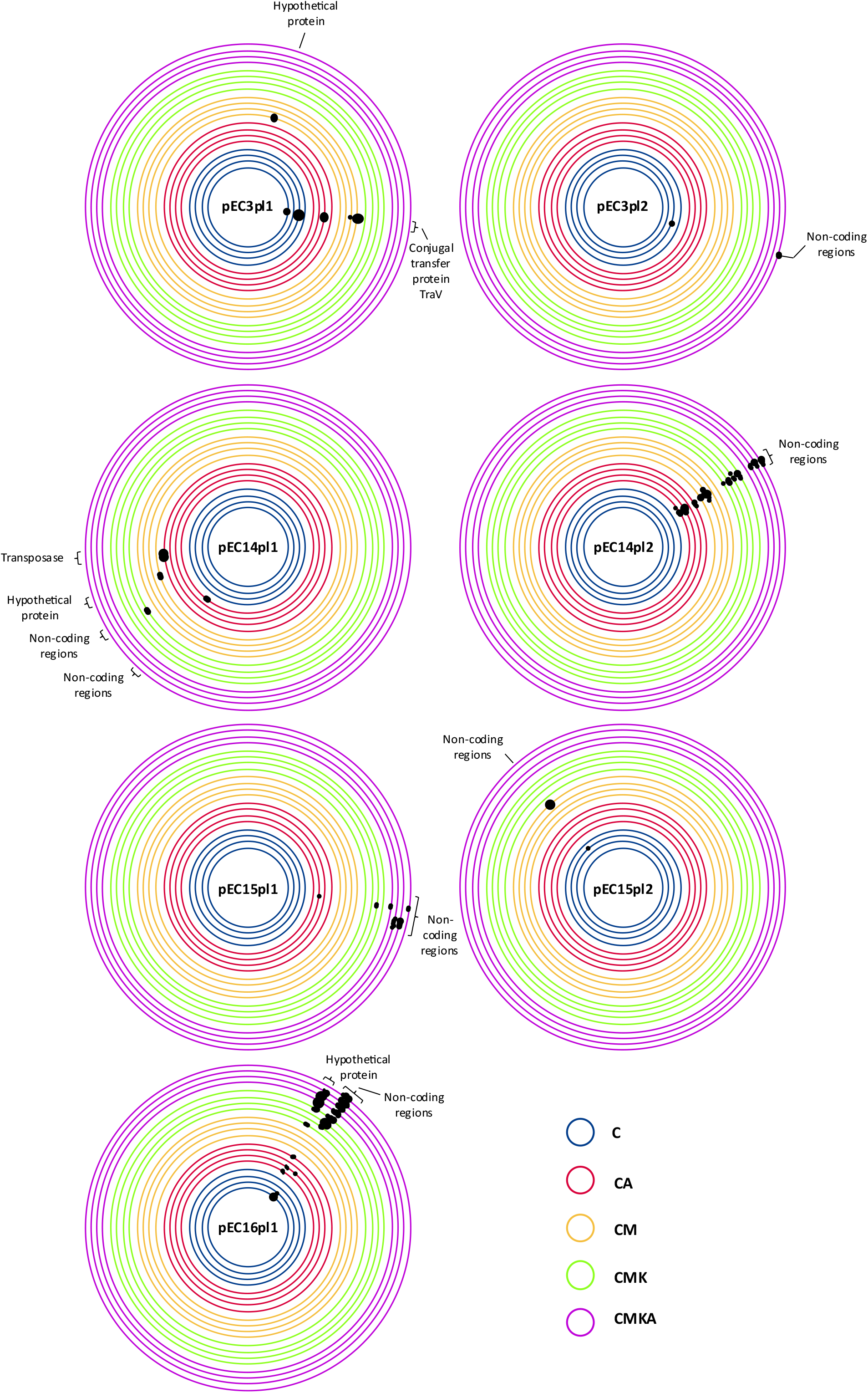
Observed mutations in plasmids after 42 cycles in the serial culture experiment (~320 generations). Only plasmids with mutations are shown. The black dots mark the mutation region and its frequency; the bigger the black dot, the higher mutation frequency (for exact values, see Supplementary data 1).

**Figure 6.**
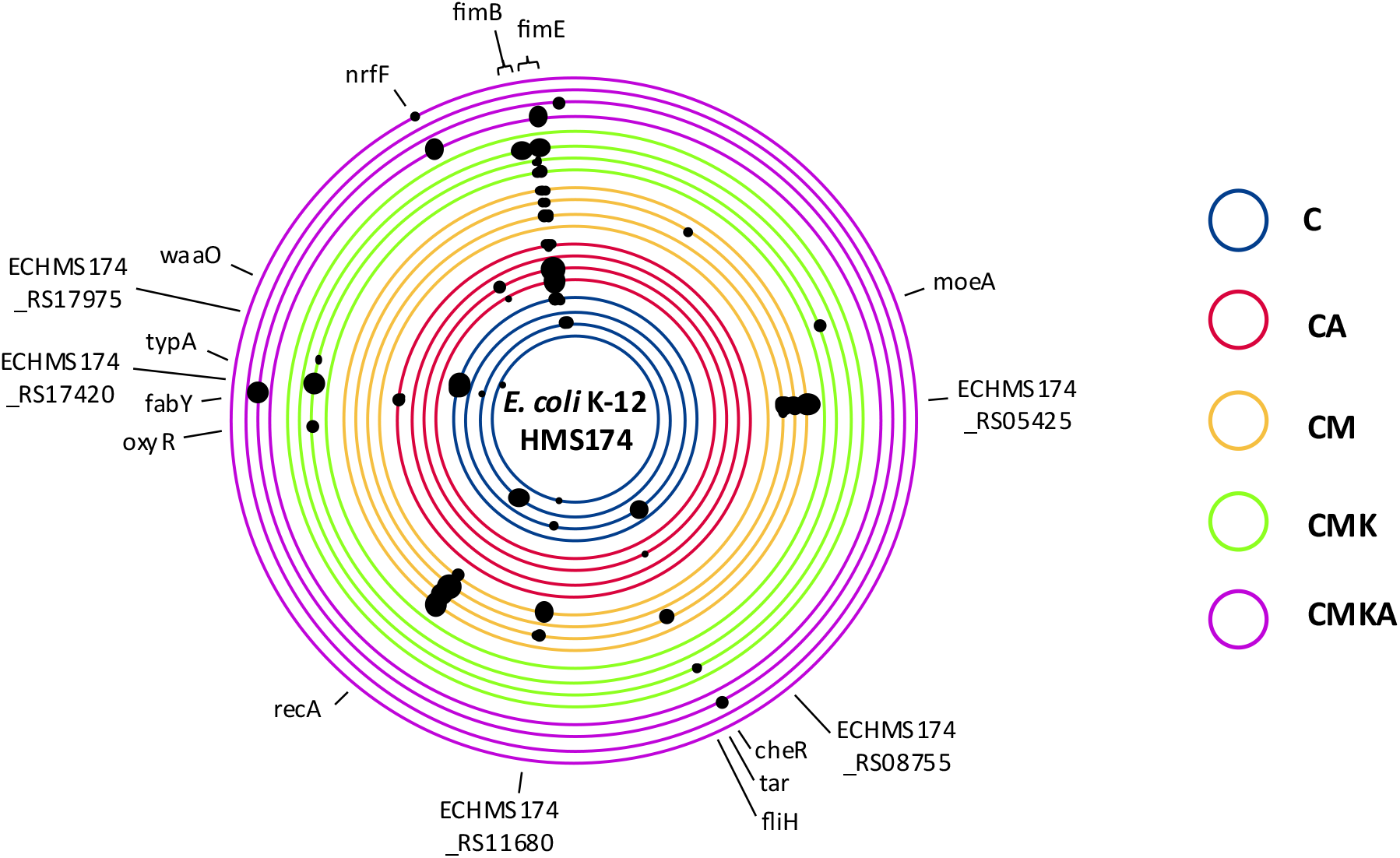
Observed mutations in the host chromosomes obtained from the community after 42 cycles in the serial culture experiment (~320 generations). Mutations in non-coding regions are not labelled. The black dots mark the mutation region and its frequency; the bigger the black dot, the higher mutation frequency (see Supplementary data 1).

## Discussion

Plasmids are ubiquitous genetic elements of bacteria. They are responsible for several notable phenotypes of their hosts, such as antibiotic resistance or increased virulence. Arguably, these opportunistic genes that provide immense benefits in only specific conditions are likely to accumulate to horizontally transferring elements. This is because genes in mobile elements are most likely to find their way to the high-fitness hosts of the community (Carattoli, 2013; Jalasvuori and Koonin, 2015; Jordt et al., 2020). Nonetheless, not all plasmids have clearly opportunistic genes (Gama et al., 2018) and plasmids co-exist in different ecological settings where only certain plasmid-backbones may find success. The potential effects that these conditions have on the survivability or dominance of plasmids in the presence of multiple competitors, to our knowledge, have not been studied directly. We set to investigate how plasmids thrive in the presence and absence of selection for opportunistic genes (antibiotic resistance) when there is or is not a migrating plasmid-free host with higher or equal fitness available. The studied plasmids represent various incompatibility types, mobility groups and conjugation systems and hence resemble a situation where different plasmids clash in a shared community.

Certain plasmids were unable to survive regardless of the tested setup. Plasmid pEC14pl1 disappeared in all experiments. It is by far the largest plasmid (143kb) and hence a notable burden to the host in the absence of selection for one of its nine resistance genes. This supports the line of reasoning where those plasmids that have accumulated large number of opportunistic genes are more dependent on specific selection on plasmid-carried genes than smaller plasmids (Ruotsalainen et al., 2020). However, and similarly to pEC14pl1, RP4 also disappeared from all communities despite of being relatively small in size (60kb). RP4 is an IncP plasmid that has a wide-host-range and it had the highest conjugation rate of all the plasmids of this experimental setup. Yet, it caused a greater reduction in the total population density compared to other plasmid combinations and therefore in a serial culture experiment, it is likely to dilute out. However, fitness cost of RP4 has been shown to readily ameliorate in *Pseudomonas* sp. host after mutations in genes for accessory helicases (Loftie-Eaton et al., 2017). Similar helicases exist in *E. coli* used in the study, but no such mutations were observed at the end of the experiment. This suggests that, while mutations may alleviate costs, the presence of alternative plasmids that are initially less costly to the host and provide similar opportunistic genes (antibiotic resistance) are likely to rapidly outcompete maladapted plasmids and hence probably reduce the chances for ameliorative mutations to establish in the community. IncP plasmids are also noted to be rare among Enterobacterial hospital pathogens (Carattoli, 2009) and hence the observed inability to survive against more common (narrow-host range) plasmids (such as IncF and IncI) is expectable. If the system had alternative hosts besides *E. coli*, the variety of potential hosts (that are generally unsuitable for the here-used pEC-plasmids) may have allowed it to prevail (Hall et al., 2016; Gama et al., 2018). All other plasmids survived in at least some conditions.

The plasmids in this study were seeded into the experiment in their natural combinations. It could be anticipated that these plasmids had already adapted to coexistence in their original hosts. However, in only some setups pEC15 and pEC16 plasmids remained at similar levels throughout the experiment (although it must be noted that our quantification method did not allow determining whether they existed in the same cells). As such, previous co-existence appears to readily dissolve and potentially new plasmid-combinations form.

Interestingly, pEC16pl2 is a small mobilizable plasmid that encodes for a (putative) microcin that is lethal to surrounding cells unless it carries the plasmid. HMS174 with pEC16 inhibits the growth of HMS174 harbouring any of the other plasmid-combinations (data not shown). Intrinsically, it appears reasonable to expect that pEC16pl2 becomes dominant in the community. Nevertheless, preliminary experiments demonstrated that pEC16pl2 does not replace other plasmids from the system even in the absence of antibiotics and, as such, we retained it in the experimental setup to evaluate the plasmid’s long-term viability. Indeed, in the absence of simulated presence of a pathogen (i.e. sub-lethal kanamycin and kanamycin resistant *E. coli*), the amount of pEC16pl2 remained among the highest. However, the introduction of kanamycin and a kanamycin-resistant host (i.e. simulated proliferating pathogen) caused it to rapidly disappear. This indicates a clear trade-off where change in the ecological setting turns the plasmid’s fitness completely around. The initially co-existing plasmid partner of pEC16p2 (i.e. pEC16pl1) was stable only in the refreshed community (C), suggesting that pEC16pl2 prevalence in other systems was independent of pEC16pl1.

Plasmids are generally divided into incompatibility groups where two plasmids of the same type cannot coexist in a single cell line. Here, plasmid pEC13 and pEC14pl3 both belong to IncFII and generally prevailed in only those cultures where the other was absent. pEC13 became the dominant IncFII plasmid in conditions where the simulated pathogen was present (CMK and CMKA) whereas pEC14pl3 prevailed in others. Correspondingly, pEC16pl1, pEC14pl2 and pEC15pl1 are IncI1 plasmids, of which pEC16pl1 was prevalent in only community C whereas genetically indistinguishable plasmids pEC14pl2 and pEC15pl1 were prevalent in others. This suggests that plasmids of the same incompatibility group with similar size and overall genetic characteristics can have drastic differences in fitness in response to the ecological condition of their hosts.

Host chromosomes are known to adapt to plasmid presence, but former studies have focused on investigating the effects of singular plasmids (see e.g. Dahlberg and Chao, 2003; Yano et al., 2016; Loftie-Eaton et al., 2017). Here in an environment with 11 interacting mobilizable or conjugative plasmids (two of which were almost identical), the only consistently occurring mutations over 300 generations were observed within gene *fimE*. This gene is responsible for switching off *fim* operon and hence fimbriae production in *E. coli* (Abraham et al., 1985). *FimE* inactivating mutations therefore result in continuously active fimbriae operon. Fimbriae play key roles in *E. coli* pathogenicity, host immune responses and biofilm production (Decano et al., 2021). IncI1 and F-plasmids plasmids have been shown to encode pili that induce fimbriae-like phenotype in the host and pilin mutants have significantly lower conjugation rate compared to wild type (Dudley et al., 2006; Virolle et al., 2020). It is possible that *fimE* inactivation and hence constitutive fimbriae production is an adaptation to the general experimental conditions where the bacteria were cultivated. However, this is unlikely to be the case since evolution occurred outside of the host organism (no benefit for pathogenicity) and attachment to surfaces (Kallas et al., 2020) would have made cells less likely to be transferred to the fresh medium in the experimental setup where planktonic population was utilized to seed the next culture cycle. Given that majority of the plasmids here encode pili with (likely) comparable phenotypes to fimbriae, it is within reason to expect that the expression of chromosomally encoded fimbriae was under selection solely due to the large number of genetically different plasmids. In other words, the generation of fimbriae-like phenotype especially from chromosome instead of one of the several potential plasmids had a specific benefit for the host bacterium. Porse and colleagues (Porse et al., 2016) observed *fimE* mutation in a serial culture experiment with a single IncF-plasmid in only one of several evolved *E. coli* lineages. In the famous long-term evolution experiment, no such mutations were observed after 2000 to 20000 generations (Barrick et al., 2009). In both of these experiments, the culture conditions were roughly the same (i.e. 37°C temperature and shaking). Nevertheless, while the detailed study of this was beyond our resources, we cautiously hypothesize that those bacterial cells that produced more fimbriae over their contemporaries may be less prone to the invading plasmids that were present in the surrounding bacteria. These other bacteria may have encoded fimbriae-like pili mainly from plasmids. To clarify, fimbriae or pili were present in most cells nevertheless and there was a difference whether it was pili that initiates conjugation or if it was fimbriae encoded by the chromosome (that do not lead to conjugation). This chromosomal expression may have protected the bacterium from plasmid invasion and hence from the recently demonstrated acquisition-associated detrimental fitness effects (Prensky et al., 2021). Therefore, this selection might be observable only in conditions where several genetically different plasmids can simultaneously enter the host and no specific mutation in any of the plasmids or in the chromosome can shut them all down.

Overall, the study shows that plasmids have trade-offs that allow them to outcompete their contemporaries only in certain conditions. The big multi-resistance-providing plasmids may require specific selection to remain viable in the presence of smaller competitors. This observation can be of importance, given that limiting the administration of multiple types of antibiotics reduces the selection for large plasmids like pEC14pl1 and hence may lead to their displacement with smaller plasmids with fewer resistance genes. On the other hand, broad host-range plasmids (like RP4) may serve as reservoirs of antibiotic resistance genes in non-pathogenic hosts in different environments from which they can supply resistances to relevant hosts (as noted by Loftie-Eaton and colleagues; Loftie-Eaton et al., 2017). In general, the absence of selection for plasmids via antibiotics did not have an effect on plasmid prevalence and hence removing antibiotics altogether does not seem to rapidly re-sensitize bacteria to drugs even in highly competitive situations where non-resistant, rapidly proliferating cells are present (i.e. CMK community). Multi-plasmid clashes are likely to occur in natural environments and under differing ecological settings. This work here provides one of the first overviews of the dynamics that occur during such multi-plasmid interaction and may illuminate situations where certain plasmids become over-represented over their contemporaries.

## Materials and methods

### Bacterial strains, plasmids and culture conditions

Bacterial strains and their plasmids are listed in Table 1. For initiating the community experiment, each strain was grown separately in 50 ml tubes containing 5 ml of Luria-Bertani broth (LB) (Bertani, 1951) with the appropriate antibiotic selection (either 150 μg/ml ampicillin or 25 μg/ml kanamycin) for overnight at 37°C with shaking 200 rpm. Due to experimental design, kanamycin gene had to be inactivated in plasmid RP4. This was done as described in Ruotsalainen et al., 2020. Briefly, a synthetic gene containing an inactivating mutation within kanamycin resistance gene was inserted with homological recombination in the RP4 plasmid using Red/ET Recombination (Gene Bridges) according to manufacturer’s protocol. Kanamycin-sensitive phenotype was screened from the obtained colonies and the presence of the inactivating insertion verified with polymerase chain reaction. The mutated plasmid was transferred via conjugation to HMS174.

### Plasmid competition experiment

Microcosm experiment was set up into five ecological settings (treatments) and named as C, CA, CM, CMK and CMKA. At the start of the experiment (cycle 1; C1), 5 μl overnight (o/n) cultures of pEC strains and RP4 were added to a 50 ml tube containing 5 ml LB medium supplemented with appropriate antibiotics and/or bacterial culture according to the setup (see Table 5 and Figure 1). The cultures were grown at 37°C and aerated by slow agitating at 60 rpm for 24 hours. The experiment was done with four identical replicates/treatment. After each cycle, 50 μl of the culture (1% inoculation) was transferred to the fresh media, and in case of CM, CMK and CMKA, 5 ul of overnight cultivated migrant added. The cultures were serially propagated for 42 cycles.

**Table 5.** Treatments and their conditions

| **Treatment** |  | **Media** |
| --- | --- | --- |
| **C** | Plasmid community without selection | 5 ml LB medium |
| **CA** | Plasmid community under lethal β-lactam selection | 5 ml LB + 150 μg/ml ampicillin |
| **CM** | Plasmid community with a constant migration of plasmid-free hosts | 5 ml LB + 5 μl plasmid-free HMS174 o/n culture |
| **CMK** | Plasmid community with a constant migration of higher fitness plasmid-free hosts | 5 ml LB + 2.5 μg/ml kanamycin + 5 μl HMS174 (pET24) o/n culture |
| **CMKA** | Plasmid community with a constant migration of higher fitness plasmid-free hosts under lethal β-lactam selection | 5 ml LB + 150 μg/ml ampicillin + 2.5 μg/ml kanamycin + 5 μl HMS174 (pET24) o/n culture |

### Sample collections and DNA extraction

The samples were harvested after every 3 cycles in the first week and then, after every 7 cycles. One ml of each bacterial culture (in total 20 cultures; 5 setups x 4 biological replicates) was collected in 1.5 ml tubes and stored at −80°C for the DNA isolation. Additionally, the cultures were collected for the measuring of growth rate and yield. Here 1 ml of each bacterial culture was mixed with 300 μl of sterile 87% glycerol in cryotubes and stored in −80°C.

The total DNA (genomic and plasmid) of all the samples from cycle 1, 14, 32, 35 and 42 were extracted with Wizard^®^ Genomic DNA purification kit (Promega) following the manufacturer’s instructions. Qubit 3.0 fluorometer was used to determine the concentration of DNA using the dsDNA high sensitivity kit (Invitrogen, ThermoFisher Scientific).

### Quantitative PCR (qPCR) with plasmid specific primers

The amount of each plasmid in the samples obtained from the community experiment was quantified with quantitative PCR (qPCR). Prior to this, each primer pair used in this experiment were optimized for their efficiency (Supplementary data 1). The crosscheck test with all other plasmids in this experiment was also performed to ensure the specificity (Supplementary data 1). qPCR was performed in triplicates for each sample. Final reaction volume of 20 μl contained 1x SsoAdvanced universal SYBR green supermix (Bio-Rad), 0.5 μM of forward and reverse primers and 2 μl of DNA template (1 ng/μl). The qPCR cycle program consisted of initial denaturation at 98°C for 3 minutes, followed by 40 cycles of denaturation at 98°C for 15 seconds and primer annealing - extension at 60°C or 65°C (depends on the primer pairs; Supplementary data 1) for 1 minute 15 seconds. All qPCR assays were performed in Bio-Rad CFX96 Touch™ Real-Time PCR Detection System (Bio-Rad).

The amount of each plasmid at every time point (cycle 1, 14, 32, 35 and 42) for all treatments and all replicates was analyzed with CFX Maestro™ 1.1 software (version 4.1.2433.1219) by normalization against 16S rRNA using normalized expression mode from gene study function.

### Bacterial growth rate and yield

A single colony of each original bacterial strain was inoculated into 5 ml media containing appropriate antibiotics: LB medium without antibiotics for plasmid-free HMS174, LB medium supplemented with 150 μg/ml ampicillin for HMS174 carrying pEC plasmids and RP4, and LB medium supplemented with 25 μg/ml kanamycin for HMS174 carrying pET24. Cultures were grown at 37°C, 200 rpm, overnight. Prior to the measurement of growth rate (see below), 50 μl of each culture was transferred into 5 ml of new media and mixed thoroughly. 200 μl of this inoculated media was transferred onto a honeycomb plate with four replicates/sample. All the strains were tested for growth under four different LB-based media: without antibiotics, with 150 μg/ml ampicillin, with 2.5 μg/ml kanamycin, and with 150 μg/ml ampicillin and 2.5 μg/ml kanamycin.

Additionally, the growth rate and yield of bacterial communities at the beginning (cycle 1) and in the end of the experiment (cycle 42) from each treatment were measured. 50 μl of thawed sample was transferred into 5 ml of the media composition used in the serial culture experiment (Table 4). From this, 200 μl was transferred into a honeycomb plate with four replicates/sample. The growth curves and maximum absorbance at 600 nm was measured with Bioscreen C MBR machine (Bioscreen, Oy Growth Curves Ab Ltd).

The maximum growth rates and yields of each strain were determined with Bioscreen C MBR machine (Bioscreen, Oy Growth Curves Ab Ltd) at 37°C with low shaking. The absorbance was measured at OD_600_ for 16 hours in 5-minute intervals. The maximum growth rate and yield were calculated from the data obtained from Bioscreen using Rstudio (version 1.1.456).

### Conjugation rate test

HMS174 (pET24) was used as a recipient strain and all the strains with pEC plasmids and RP4 were the donor strains. All the strains were sub-cultured from frozen stock prior to the test.

A single colony of recipient strain was inoculated into 50 ml falcon tube containing 5 ml LB medium supplemented with 25 μg/ml kanamycin whereas each of the donor strains was grown separately in 5 ml LB medium supplemented with 150 μg/ml ampicillin. All the cultures were grown at 37°C overnight, 200 rpm. Each one of the donors and recipient strain, a total of 6 different combinations, were then mixed at 1:1 volume ratio (5 μl each) in a 50 ml falcon tube containing 5 ml LB medium without antibiotics. The test was done in four biological replicates/combination. The cell mixtures were then incubated at 60 rpm, 37°C for 24 hours. After incubation, the cultures were serially diluted to the appropriate dilutions (from 10^-2^ to 10^-6^) and plated on LB agar with 150 μg/ml ampicillin and 25 μg/ml kanamycin. The plates were incubated overnight at 37°C. Each strain without the conjugation was also plated as the negative control. The conjugation rate was calculated as colony-forming unit/ml (CFU/ml).

### Mutation mapping

The DNA samples from the end-point of the experiment were used for preparation of sequencing library with NEB Next^®^ Ultra™ DNA Library Prep Kit (Cat No. E7370L) and sequenced on Illumina NovaSeq 6000 platform with S4 flowcell (PE150).

After receiving the sequence data, the mapping of plasmid sequences to find possible mutations and to compare between treatments was performed with CLC Genomic Workbench software version 11 (Qiagen). First, the reference genomes of *E. coli* HMS174 and all the pEC plasmids, RP4 and pET24 were combined as a single reference file using Geneious Prime software version 2020.1.2 (Geneious). Then, the reference genomes and Illumina sequence reads of all the samples were imported into the workbench. The reads were then mapped to the references, followed by variant detection with 35% as a threshold for mutation and annotation. Finally, the data was exported to Microsoft Excel for further analysis.

### Statistical analysis

All statistical analyses were performed using SPSS (version 26, IBM SPSS). Two-way analysis of variance (ANOVA) with Tukey HSD as post hoc was used to analyze the difference in growth rate and yield between strains and between communities under all treatments at the beginning and the end of the experiment. The conjugation rate was analyzed with one-way ANOVA, post hoc = Tukey HSD. P-values <0.05 were considered significant in all tests.

## Notes

### Competing Interest Statement

The authors have declared no competing interest.

